# Clonal expansion dictates the efficacy of mitochondrial lineage tracing in single cells

**DOI:** 10.1101/2024.05.15.594338

**Authors:** Xin Wang, Kun Wang, Weixing Zhang, Zhongjie Tang, Hao Zhang, Yuying Cheng, Da Zhou, Chao Zhang, Wen-Zhao Zhong, Qing Ma, Jin Xu, Zheng Hu

## Abstract

Although mitochondrial DNA (mtDNA) variants hold promise as endogenous barcodes for tracking human cell lineages, their efficacy as reliable lineage markers is hindered by the complex dynamics of mtDNA in somatic tissues. Here, we utilized computational modeling and single-cell genomics to thoroughly interrogate the origin and clonal dynamics of mtDNA lineage markers across various biological settings. Our findings revealed the majority of purported “clone informative variants (CIVs)” were pre-existing heteroplamies in the first cell instead of *de novo* somatic mutations during divisions. Moreover, CIVs demonstrated limited discriminatory power among different lineages during normal development; however, certain CIVs with consistently high allele frequencies proved capable of faithfully labeling cell lineages in scenarios of stringent clonal expansion. Inspired by our simulations, we introduced a lineage informative (LI) score, facilitating the identification of reliable mitochondrial lineage markers across different modalities of single-cell genomic data.

## Main

Mitochondrial DNA (mtDNA) mutations naturally occur and accumulate over cell divisions, thus exhibiting promises as a natural barcode for tracking human cell lineages. Given its advantage in inferring clonal relationships in native human tissues, mtDNA mutations have been utilized to study the clonal dynamics underlying normal development and disease^1–4^. Rapid development of methods for simultaneous single-cell profiling of mtDNA mutations and chromatin accessibility (e.g. mtscATAC-seq^4,5^) or transcriptome (e.g. MAESTER^6^) or both (e.g. ReDeeM^2^) has greatly accelerated and expanded the application of mitochondrial lineage tracing technologies^7^. However, the complex dynamics of mitochondrial replication over cell divisions, including mitochondrial bottleneck and random segregation of mitochondrial DNA into daughter cells^8,9^, introduces a high degree of dynamics and complicates the use of mtDNA mutations as clone-informative markers. Therefore, it necessitates a systematic study on characterizing the genuine lineage-informative mtDNA mutations.

Here, we developed a computational framework to simultaneously simulate mtDNA mutations and single-cell transcriptomes under two distinct biological scenarios: clonal maintenance and clonal expansion, resembling normal tissue renewal and somatic evolution (e.g. cancer), respectively. Our results showed that a significant proportion of the identified CIVs was already present (pre-existing) in the initial cell, which is a distinct feature of mDNA unlike nuclear genomic variants-based lineage tracing. More importantly, CIVs identified using current method demonstrated poor performance in reconstructing true cell lineage history especially under scenarios of normal tissue development. However, certain mtDNA mutations with average of high VAF but low variance could still be informative for labeling specific lineages under strong clonal expansions. Overall, our study interrogated the dynamics of mtDNA mutations and demonstrated scenarios with significant clonal expansions as optimal biological settings for applying mitochondrial lineage tracing methods.

To better understand the dynamics of mtDNA mutations, we developed a computational framework to simulate the replication and random segregation of mitochondrial genomes in a cell population over division and differentiation (**Fig. 1A, Supplementary Fig. 1A-B, Methods**). Briefly, mitochondrial genomes in the first cell were initialized with pre-existing mutations where the initial heteroplasmy levels (variant allele frequency or VAF) followed a power-law distribution^10^ (**Supplementary Fig. 1C**). The mitochondrial genomes accumulated *de novo* mutations at a specific rate during replication (*μ* = 0.0008 per replication per mitochondrial genome^11^), while all mitochondrial genomes, regardless of whether they are newly generated or not, were randomly segregated into two daughter cells during division (total mutation rate per cell division *u* = 0.0008 × 500 = 0.4, when assuming 500 mtDNA copies per cell^12^). To interrogate the effects of mitochondrial bottleneck, a process in which the mtDNA copy number within a cell dramatically decreases^9^, we designed two models wherein mtDNA copy number sharply drops and resumes (referred as MT-bottleneck model hereafter) or maintain constant (referred as MT-constant model hereafter, **Supplementary Fig. 1A-D**). In parallel with the cell division, we simulated transcriptomic profiles (n = 4,000 genes) for each cell and designated cell states according to linear or bifurcated differentiation patterns using our previously developed simulation framework^13^ (**Supplementary Fig. 1A-B, Methods**). Two scenarios of clonal dynamics, including clonal maintenance and clonal expansion (**Fig. 1B, Supplementary Fig. 1A-B**), were further simulated to assess the performance of mitochondrial lineage tracing. Clonal maintenance corresponds to normal developmental processes, where most cell lineages remain over stem cell self-renewal. In contrast, clonal expansion corresponds to lineage turnover commonly observed in cancer progression, immune responses, *etc*.

**Fig. 1.**
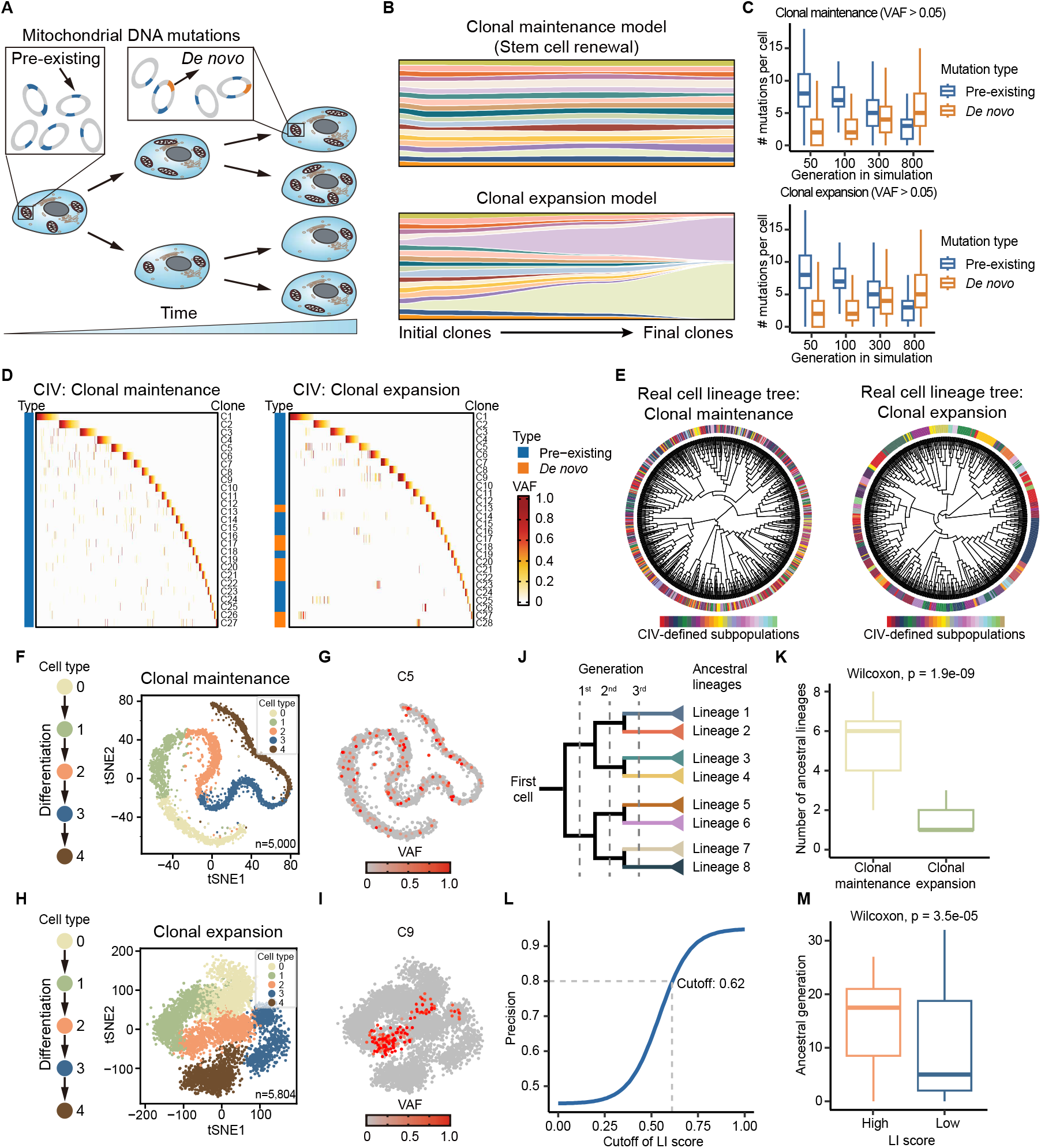
Computational simulation of mitochondrial lineage tracing in single cells. **A**, Schematic of mitochondrial lineage tracing. Mitochondrial genomes that carry pre-existing mutations (blue) in the initial cell were replicated and segregated into daughter cells, and *de novo* mutations (orange) continuously arose during cell divisions. **B**, Schematic of clonal maintenance (lop) and clonal expansion (bottom) model, respectively. **C**, Number of pre-existing (blue) or *de novo* (orange) mutations per cell over cell divisions in clonal maintenance (top) or clonal expansion model (bottom). **D**, Identification of clone informative variants (CIVs) in clonal maintenance (left) or clonal expansion model (right). Heatmaps showing the variant allele frequencies (VAFs) of CIVs in single cells. Rows represents CIVs and columns represent single cells. **E**, Ground-truth cell lineage tree with CIV-defined subpopulations annotated on the tree. **F**, Diagram of I-distributed stochastic neighbor embedding (tSNE) for simulated transcriptomic profiles of single cells, following linear differentiation (left) in clonal maintenance model. **G**, An exemplar CIV (C5) where the defined subpopulation of this variant is highlighted. **H**, ISNE diagram of simulated transcriptomic profiles of single cells, following linear differentiation (left) in clonal expansion model. **I**, An exemplar CIV (C9) where the defined subpopulation of this variant is highlighted. **J**, The eight ancestral lineages of 3rd generation of each cell were traced to analyze the lineage composition of CIV-defined subpopulations. **K**, Comparison of ancestral lineages of each CIV-defined subpopulation between clonal maintenance model and clonal expansion model. **L**, Line graph showing the precision of CIV varies with different lineage-informative (LI) score cutoffs. The precision reaches 0.8 when LI score cutoff is set to 0.62. **M**, Comparison of ancestral generations of CIV-defined subpopulations between high LI score(≥ 0.6) groups and low LI score(< 0.6) groups.

First, we checked the mutation dynamics over the simulations and found that the number of pre-existing mutations per cell constantly decreased while *de novo* mutations gradually accumulated in both clonal maintenance and clonal expansion scenarios (**Fig. 1C**). This is expected because a pre-existing heteroplasmy can be lost in descendant cells due to mitochondrial segregation and random drift^14^. A CIV is defined by its unique presence in a cell subpopulation, which is regarded as a unique marker for a cell clone^6^. Indeed, CIVs from both models can be identified using the method commonly used in the literature^6^ (**Fig. 1D, Methods**). In fact, CIVs in clonal maintenance model were mostly pre-existing mutations even after 800 cell divisions (> 99%, **Supplementary Fig. 2A**), whereas some CIVs in clonal expansion model were indeed *de novo* mutations (10.7%, **Supplementary Fig. 2B**), indicating *de novo* MT mutations expanded along with clonally expanded cells. Given the availability of genuine cell division history in our simulations, lineage relationship of CIV-defined subpopulations could be assessed. Surprisingly, we observed that cells within each CIV-defined subpopulation failed to aggregate in the ground-truth cell lineage tree in clonal maintenance model (**Fig. 1E**). Interestingly, clonal expansion resulted in a prominent pattern of phylogenetic aggregation for CIV-defined cell subpopulations, indicating high efficacy of these CIVs in distinguishing different cell lineages. We then examined the similarity of transcriptomes of cells within each CIV-defined subpopulation for both models (**Fig. 1F-I**) and we found that certain CIV-defined subpopulations (e.g. C9) in clonal expansion model showed a high degree of transcriptomic similarity and displayed clustered patterns on the low dimensional transcriptomic space through t-distributed stochastic neighbor embedding (t-SNE) (**Fig. 1I, Supplementary Fig. 2C**). On the contrary, clonal maintenance model failed to show such clustered patterns where each CIV-defined subpopulation consists of almost all cell types modeled in simulation (**Fig. 1G, Supplementary Fig. 2C**).

To further quantify the lineage tracing capacity of CIVs, we defined cells in the third generation of divisions from a common progenitor cell as lineage ancestors (termed Lineage 1-8, **Fig. 1J**) and traced the origin of each CIV-defined subpopulation back to these eight lineages (**Methods**). In line with aforementioned findings, CIV-defined subpopulations in clonal maintenance model each consisted of a large number of ancestral lineages (median = 6 out of 8), thus displaying a highly mixed linage composition. However, clonal expansion model showed a low number of ancestral lineages (median = 1 out of 8), suggesting better performance of CIVs in the context of clonal expansion (**Fig. 1K**). To comprehensively assess the efficacy of mitochondrial lineage tracing in more diverse contexts, we conducted the same analysis for MT constant model (without mitochondrial bottleneck, **Supplementary Fig. 3**) and bifurcated differentiation (**Supplementary Fig. 4** and **5**), and the results were consistent with MT bottleneck model under linear differentiation as shown in **Fig. 1**. Collectively, these results demonstrated that clonal expansion is a prerequisite for applying mitochondrial lineage tracing methods.

Inspired by these findings from simulations, we next sought to introduce a quantitative metric for identifying the reliable lineage-informative mtDNA variants in single-cell genomic data. Clearly, if an mtDNA variant is fixed (VAF∼100%) in a cell subpopulation (referred as subclonal homoplasmy), this variant stably labels the corresponding cell lineage over time, just as a variant in nuclear genome. However, subclonal homoplasmies are rare because most subclonal variants are heteroplasmy within a cell. We therefore defined a metric to quantify the reliability of CIV as reliable lineage marker,

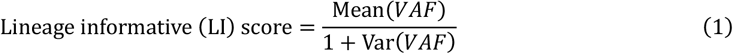

where Mean(VAF) and Var(VAF) denote the average and variance of VAFs, respectively, observed in the cells with the detected variant. A higher LI score indicates higher reliability of this mtDNA variant as cell lineage marker. To establish a practical threshold for the LI score, we generated a precision-versus-cutoff curve (**Methods**). Our analysis revealed that setting the LI score cutoff at approximately 0.6 (0.62) achieved an 80% precision rate in our simulation data. (**Fig. 1L**). Above this cutoff, there was a strong diminishing return for the increase of precision rate. We then adopted the cutoff of 0.6 and compared the ancestral generation of each CIV-defined subpopulation with high LI scores vs with low LI scores. Notably, the ancestral generation of CIV-defined subpopulation with a high LI score was significantly later than that with low LI score (**Fig. 1M, Supplementary Figs. 3J, 4J, 5J**), suggesting that this cutoff indeed could be used for categorizing CIVs. Hence LI score ≥0.6 was used to select good CIVs in real single-cell genomic data analysis.

We first analyzed a real single-cell RNA-seq dataset (Smart-seq2) derived from an early human embryo^15^ (**Fig. 2A**). The high coverage (∼50x for each cell) of mitochondrial genome enabled the detection of mtDNA mutations at single-cell resolution (**Supplementary Fig. 6, Methods**). Following that, eight CIVs were identified using the method in the literature^6^ (**Fig. 2B**). Consistent with the simulation results of clonal maintenance, although CIVs could be identified, their LI scores were generally low (7 out of 8 had LI scores<0.6) and only one mtDNA variant (15959G>A, LI score=0.74) passed our CIV selection (**Fig. 2B**). The cell type compositions of these CIV-defined subpopulations, including 15959G>A, were also highly diverse (**Fig. 2C-D**). These data together indicate a poor performance of mitochondrial lineage tracing in contexts with no significant clonal expansions, such as embryonic development.

**Fig. 2.**
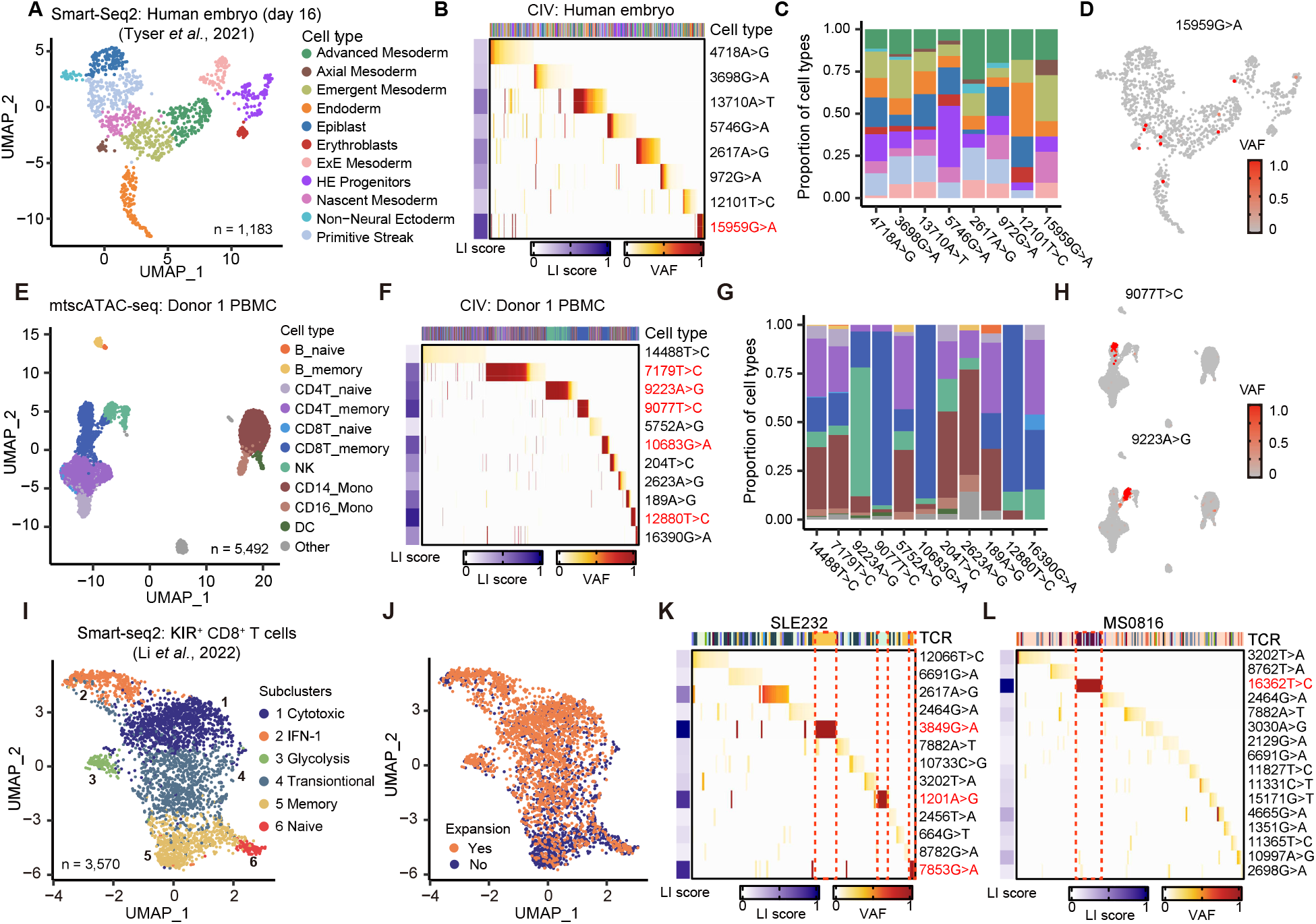
Mitochondrial lineage tracing in real single-cell genomic datasets. **A**, Single-cell transcriptomic profiles of an early human embryo (day 16) generated by Smart-seq2. **B**, Heatmap showing CIVs identified in this early embryo dataset. CIVs with LI score ≥ 0.6 are highlighted. **C**, Cell type composition of each CIV-defined subpopulation. **D**, Distribution of 15959G>A-defined subpopulation on UMAP diagram of transcriptomic profiles. **E**, Single-cell chromatin accessibility profiles of PBMCs from an old individual (73 years old), Donor 1. **F**, Heatmap showing the identification of CIVs in Donor 1. **G**, Cell type composition of each CIV-defined subpopulation in Donor 1. CIVs with LI score ≥ 0.6 are highlighted. **H**, Distribution of 9077T>C- and 9223A>G-defined subpopulations on the single-cell chromatin accessibility profile, respectively. **I**, Single-cell transcriptomic profiles, generated with Smart-seq2, of KIR+ CD8+ T cells from healthy individuals and autoimmune diseases. **J**, Clonal expansion status of profiled T cells, determined by TCR sequence. **K**, Heatmap showing CIVs in a systemic lupus erythematosus (SLE) sample - SLE232. Red dotted rectangles highlight cell subpopulations defined by 3849G>A, 1201A>G and 7853G>A with high LI scores (≥ 0.6). **L**, Heatmap showing CIVs in a multiple sclerosis (MS) sample - MS0816. The red dotted rectangle highlights the cell subpopulation defined by 16362T>C (LI score ≥ 0.6).

For the validation of the results of clonal expansion model, we collected peripheral blood mononuclear cells (PBMC) samples from two donors (one male and one female) over 70 years old because clonal hematopoiesis, a typical clonal expansion process, frequently occurs in elderly individuals^16^. We performed mtscATC-seq of PBMC samples from these two individuals and the cell types were annotated using chromatin accessibility of single cells (**Fig. 2E, Supplementary Fig. 7**). With an average of 20-40X coverage of mitochondrial genome per cell, we identified mtDNA mutations and subsequently performed CIV analysis in both samples (**Fig. 2F, Supplementary Fig. 7**). We found that 5 out of 11 mtDNA variants can be selected as good CIVs (LI score ≥ 0.6). Interestingly, compared with CIVs with low LI scores, reliable CIVs showed a biased cell type composition. For example, CD8^+^ T memory cells made up 89.3% of cell subpopulations defined by 9077T>C and NK cells made up 66.0% of cell subpopulations defined by 9223A>G (**Fig. 2G**). Furthermore, cells within the 9077T>C or 9223A>G subpopulation exhibited a significantly closer relationship in terms of transcriptomes, indicating that those CD8^+^ T memory cells or NK cells possibly arose from a common progenitor cell by clonal expansions (**Fig. 2H**). Analysis of another independent sample, Donor 2, showed similar results as Donor 1 (**Supplementary Fig. 7**). Overall, these results, unlike the human embryo data (**Fig. 2A-D**), strongly suggested that CIVs identified in the context of clonal expansion, particularly those with high LI scores, were promising lineage markers, further supporting our simulation results.

Although these results agreed with the simulations, using transcriptomic similarity alone for testing mitochondrial lineage tracing could lead to false conclusions due to the inconsistency between transcriptomic similarity and lineage relationship under certain circumstances of lineage plasticity. Therefore, we took advantage of another Smart-seq2 dataset including T cells isolated from patients with autoimmune diseases (**Fig. 2I**)^17^. TCR (T-cell receptor) sequence is expanded along with the expansion of T cells under immune responses, thus serving as a reliable lineage marker independent of mtDNA mutations. We then sought to use TCR-defined clonal structure as ground truth to evaluate the efficacy of mitochondrial lineage tracing in immune responses with significant clonal expansions (**Fig. 2J**). We first performed mtDNA mutation calling in sample SLE232, a patient with systemic lupus erythematosus, and MS0816, a patient with multiple sclerosis (**Supplementary Fig. 8A-D**). Identification of CIVs in both samples showed several CIVs with high heteroplasmy (both average VAF and LI score > 80%), which included 3849G>A, 1201A>G and 7853G>A in SLE232 and 16362T>C in MS0816. Importantly, we compared the subpopulations defined by CIVs and TCR sequences and found a high concordance between these CIV-defined subpopulations and TCR-defined subpopulations (**Fig. 2K-L, Supplementary Fig. 8E-H**), suggesting that these CIVs with high LI scores could be used to identify real clones. It was also worth noting that only these four CIVs (3849G>A, 1201A>G, 7853G>A and 16362T>C) showed good potential for lineage tracing, again indicating a large amount of purported CIVs are not faithful lineage markers. Together, our single-cell genomics data analysis and simulations demonstrated that clonal expansion dictates the performance of mitochondrial lineage tracing at single cell level.

In summary, our integrated study with computational simulations and real single-cell genomic data analysis demonstrated that the efficacy of mitochondrial lineage tracing was highly context-dependent. Specifically, vast majority of mtDNA mutations have limited efficacy to reconstruct lineage history in contexts of no significant clonal expansions as they failed to distinguish clonal lineages. However, they did show promises in tracing cell lineages in scenarios with strong clonal expansion as demonstrated by our simulations and human PBMCs and TCR datasets. Therefore, applications of mitochondrial lineage tracing techniques should consider scenarios undergoing significant clonal expansion. For example, tumor relapse following treatment (e.g. chemotherapy) is often accompanied by clonal expansion of a subgroup of cells with higher fitness, representing a suitable setting for mitochondrial lineage tracing^3^. Importantly, our study provided a novel method (LI score) for identifying reliable mtDNA barcodes for lineage tracing. Application of this method to emerging datasets generated by mitochondrial-enriched single-cell sequencing methods, e.g. mtscATAC-seq^4,5^, MAESTER^6^ and ReDeeM^2^, will facilitate biological discoveries. Therefore, future work that aims to obtain high sequencing coverage of mitochondrial genome alongside multi-modal information^2^, including spatial location, transcriptome and epigenome, would bring novel insights about mitochondrial lineage tracing as well as deepen our understanding of cell-fate decisions and clonal dynamics in human somatic cells.

## Methods

### Simulation of mitochondrial lineage tracing and single-cell transcriptomic data

The simulation of mitochondrial lineage tracing data is divided into three parts, simulation of stem cells growth, simulation of mitochondrial genome proliferation along with cell division and simulation of single-cell transcriptomic data.

#### Simulation of stem cells growth

As our previous published study^13^, we first simulate the stem cells growth using Gillespie algorithm^18^. By conceptualizing cell division as components of continuous-time Markov process, we designate the reaction rate of cell division as *p*(*t*) given as follows:

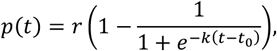

Where the three parameters *r, k* and *t*_0_ jointly determine the change of cell growth rate with time.

We then simulate the cell population growth with given division rate from 1 initial cell until the population size reached 20,000 cells. 5,000 cells were randomly sampled to obtain their division history (**Supplementary Fig. 1A**).

#### Simulation of mitochondrial genome proliferation over cell divisions

To model the mitochondrial lineage tracing within a specified phylogenetic tree, an initial population of mitochondrial DNA was allocated to the progenitor cell. The proliferation dynamics of mitochondrial DNA were simulated using the Gillespie stochastic simulation algorithm, with division and death rates at 1 and 0.1, respectively. This simulation was terminated when the mitochondrial genome count reached 500 units. During the initial phase of mitochondrial genome generation, the mutation rate per division was set to follow a Poisson distribution with an expectation of 0.2 mutations per division (**Supplementary Figure 1A, B**).

After establishing the initial population of mitochondrial DNA, the model was extended to simulate the accumulation of *de novo* mutations within the mitochondrial genome as cellular division occurred and lineages diverged. Mitochondrial DNA replication depended on cellular division, with these conditions: for mitochondrial genome doubling index *di* = 1, a single replication event for all mitochondrial DNA; for *di* within the interval (1, 2), a single replication for all mitochondrial DNA followed by a secondary replication of a random sample of *di* − 1 mitochondrial DNA; and for *di* < 1, a single replication of a randomly selected *di* mitochondrial DNA. In the simulation of MT-constant model, we set the doubling index to *di* = 1. In the simulation of MT-bottleneck model, we set *di* = 0.3 for the first 5 cell divisions and *di* = 1.2 for the latter (**Supplementary Fig. 1A, C**). When the cell divides, the replicated mitochondrial DNA are segregated to the two daughter cells, which follows binomial distribution. It has been estimated that per-mitosis mutation rate for mitochondrial genome is 10-100 higher than nuclear genome per site^19^. Assuming the somatic mutation rate of nuclear genome as ∼10^−9^ per mitosis per site^20^, 50-fold increase in mutation rate for mitochondrial genome and 500 mitochondrial copies, each cell acquires 10^−9^ × 16,569 × 50 × 500 ≈ 0.4 mitochondrial mutations per mitosis on average. Hence, we set the number of mitochondrial mutations generated within the cell after each division to follow a Poisson distribution within an expectation of 0.4 mutations per division.

To simulate the clonal maintenance model, which is the renewal of stem cells in real biology, we continued to simulate the replication and random allocation of mitochondrial DNA as cells divide. After each cell division, we select one of the two progenies to continue simulating its division process. We repeated this procedure 800 times to record the information of mitochondrial DNA mutations in different generations (**Supplementary Fig. 1A**).

To simulate the clonal expansion model, we continued to simulate the replication and random allocation of mitochondrial genomes as cells divide. Initially, all cells underwent a simultaneous division, accompanied by mitochondrial DNA replication and random allocation, after which half of the daughter cells were selected as the subsequent generation. We repeated this procedure 800 times to record the information of mitochondrial DNA mutations in different generations (**Supplementary Fig. 1A**).

#### Simulation of cell differentiation and single-cell transcriptomic data

We included simulations of cell differentiation after the stem cell growth completed. Each stem cell differentiates according to a given differentiation model (e.g. linear model or bifurcated model) to generate progenitor cells of each cell type. The division of progenitors are in asymmetric division. When a progenitor cell divides, one of its two daughter cell is still a progenitor cell that can self-renew and differentiate, and the other is a differentiated cell that no longer self-renew (**Supplementary Fig. 1A**). Cell differentiation was also simulated with a continuous-time Markov process. There is a given probability *q*_*ij*_(*t*) = 1 − *p*_*i*_(*t*) that cell type *i* could differentiate to cell type *j* at each cell division. With the cell division history and cell types generated in the previous step, we can simulate the gene expression data of different cell types following previously published study^21^. For each cell, we simulated 4,000 genes, 40% of which were cell-type specific, and these genes followed their respective expression programs within each type of cell, other genes follow the same expression program in all cell types. We assume that there are two steps in gene expression. First, we generate a time-dependent latent expression *z*_*t*_ that follows a normal distribution,

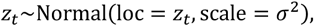

where *t* is the generation of cell. After obtaining the latent expression quantity of all genes, we sample the expression of each generation following a ZINB distribution,

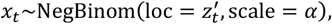

where 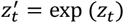, *α* is the scale parameter.

### mtscATAC-seq of PBMCs

Peripheral blood samples from two elderly individuals (Donor 1: male, 73 years old; Donor 2: female, 79 years old) were collected from Guangdong Provincial People’s Hospital. The study protocol was approved by the local ethics committee, and written informed consent was obtained from all donors. The blood samples were first diluted 1:1.5 in 0.9% physiological saline and peripheral blood mononuclear cells (PBMCs) were then isolated with a Lymphoprep™ density gradient (density 1.077 g/mL) according to the manufacturer’s instructions (STEMCELL).

scATAC-seq libraries were generated using the 10X Chromium Controller and the Chromium Next GEM Single Cell ATAC kit according to the manufacturer’s instructions. To retain mtDNA, permeabilization was done using 10 mM Tris-HCl pH 7.4, 10 mM NaCl, 3 mM MgCl2, 0.1% NP40 and 1% BSA as previously described (3, 12). PBMCs were then fixed with 1% formaldehyde (28906, Thermo Fisher) in PBS for 10 minutes at room temperature, followed by quenching with 125 mM glycine solution. Fixed PBMCs were then washed twice with PBS and centrifuged at 400g for 5 min at 4 °C. Subsequently, PBMCs were lysed with lysis buffer (10 mM Tris-HCl pH 7.4, 10 mM NaCl, 3 mM MgCl2, 0.1% NP40, 1% BSA) for 3 minutes, followed by adding 1 ml of ice-cold wash buffer (10 mM Tris-HCl pH 7.4, 10 mM NaCl, 3 mM MgCl2, 1% BSA). Then lysed cells were centrifuged at 400g for 5 min at 4 °C and the supernatant was discarded. The pellet was resuspended in 1× Diluted Nuclei buffer (10X Genomics) for counting using Trypan Blue. The downstream procedure, including tagmentation, single-cell Gel Bead-In-Emulsions (GEMs) preparation and library amplification, was performed according to the standard protocol provided by 10X Genomics. The final libraries were quantified using a Qubit dsDNA HS Assay kit (Invitrogen) and the DNS fragment was analyzed using Bioanalyzer 2100 system (Agilent).

### mtscATAC-seq data preprocessing and cell type annotation

Raw reads were mapped to hg38 reference genome using cellranger-atac (2.1.0) with default settings and the resulting fragment files were subject to ArchR (1.0.2) for downstream analysis. Cells with TSS enrichment score > 5, fragment number ≥ 1000, BlacklistRatio < 0.1 and PromoterRatio > 0.1 were kept for downstream analysis. Then doublet score for each cell was calculated and potential doublets were removed. Dimensionality reduction and clustering were carried out using addIterativeLSI, addClusters and addUMAP from ArchR package. After that, markers of each cluster were identified with cutoff FDR < 0.01 and log2FC ≥ 1 and these markers were later used for curation of cell types. To better annotate the cell type, a publicly available single-cell RNA-seq dataset of PBMC^22^ was integrated with the scATAC-seq data. Based on the integration results, we then manually curated the cell type annotation using marker gene lists from ScType database^23^ and relevant literature^9^.

### Smart-seq2 data preprocessing

Raw reads of Smart-seq2 datasets were first aligned to hg38 reference genome using STAR (2.7.10a) with default parameters. Bam files were then sorted using samtools (1.15.1) and duplicates were removed using MarkDuplicates from GATK toolkit (V4.2.0.0). Resulting bam files were then used for downstream mitochondrial mutation calling.

### Identification of mitochondrial mutations

Mitochondrial DNA mutation calling was performed as previously described^9^ with minor modifications. For mtscATAC-seq data, samtools (1.15.1) was used to subset chrM-specific alignments from the original bam file generated by cellranger-atac. The resulting bam was then subject to IndelRealigner to correct potential mapping errors around indels. After realignment, mutations were called using varscan (2.4.4) with --min-var-freq 0.01 --min-reads2 2 to identify germline mutations (bulk VAF cutoff 90%). For identification of mitochondrial DNA mutations in single cells, the realigned bam file was first split into separate bam files. For each bam file, duplicates were removed and mutations were called using varscan with the same parameter for bulk sample calling. Apart from the germline mutations identified previously, mutations that are present in more than 90% of cells with VAF > 5% were also considered as germline mutations. To identify high-confidence mutations, germline mutations were firstly removed. Second, mutations within the range of 302-316, 514-524 and 3106-3110 were also removed due to a large number of homopolymers, which potentially cause misalignment. Third, mutations must meet the following criteria: 1) variant counts ≥ 4; 2) sequencing depth of the site ≥ 20; 3) VAF ≥ 5%; 4) ratio of reads mapped to forward and reverse strand must be in the range of 0.3-0.7. After the identification of high-confidence variants in each individual cell, all high-confidence variants were merged. Following that, a mutation recalling procedure was done in the cell population. More specifically, for each high-confidence variant site, VAF was calculated for all cells. At last, cells with a mean sequencing depth < 10 were discarded.

For Smart-seq2 datasets, several extra steps were done to ensure the compatibility with the previous framework. Read groups (RG) of LB and SM were added to bam files using samtools addreplacerg. Moreover, SplitNCigarReads from GATK was used to split reads that contain Ns in their cigar string, which is caused by spanning splicing events of RNA-seq data.

### Evaluation of clone informative variants

To assess the potential of mitochondrial lineage tracing, we identified clone informative variants (CIVs) from simulated data and real data to examine their efficacy in reconstructing cell lineages. CIVs were identified using the method described by Miller *et al*^6^. We first calculated the presence of each variant in the cell population under different thresholds of VAF, ranging from 0 to 50%. We then tested various thresholds of minimal VAF and minimal cell population size to select clone informative variants (CIVs). More specifically, for each variant, only cells with VAF exceeding the minimal VAF cutoff are counted and if the number of cells carrying this variant exceeds the minimal cell population size, the variant is considered as a CIV. CIVs identified under a certain combination of thresholds were then sorted by their presence in the cell population and cells were subsequently assigned to CIVs, constituting CIV-defined subpopulations. Small CIV-defined subpopulations (< 5 cells in simulation data) were excluded from downstream analysis. Cells within each CIV-defined subpopulation were also sorted by their VAF. Pearson correlation of CIVs was performed and hierarchical clustering was conducted to calculate the distance amongst CIVs. We chose 0.8 as the cutoff for height (output of the previous hierarchical clustering) and CIVs had a distance lower than 0.8 were grouped. If multiple CIVs were clustered together (co-existed in the same subgroup of cells), the CIV with the highest mean VAF was kept. It was also worth noting that mitochondrial mutational profiles of each dataset could be greatly different from each other. For example, some cell populations had a great number of mutations with high VAF (e.g. aged PBMC samples) while others only had a few such mutations, so that the threshold of VAF and cell population size for CIV identification should be customized. Therefore, in our study the parameter for identifying CIVs in each dataset was adjusted accordingly and the final size of the cell population with CIVs should be at least 10% of the examined cell population.

To better assess the lineage tracing potential of CIVs, we examined the statistical properties of CIVs within both real and simulated datasets and observed that CIVs identified high-confidence clones exhibit a higher VAF. Moreover, the VAF distribution across all cells within a given CIV-defined subpopulation demonstrated relative uniformity. Consequently, we proposed the following metric for CIV assessment:

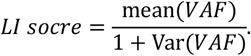

CIVs with elevated LI scores are typically indicative of clonal identification capability. To establish a cutoff LI score for clone delineation, we scrutinized simulated datasets derived from two differentiation models, linear and bifurcated, and two mitochondrial number models, constant and bottleneck. A CIV is deemed efficacious if the common ancestor of at least 60% of the cells with the closest lineage relationship within a CIV-defined subpopulation is later than 10^th^ generation. The precision of a CIV is quantified as the ratio of efficacious CIVs to the aggregate count of CIVs exceeding the cutoff score.

Analysis of the simulation data facilitated the construction of a precision-versus-cutoff curve. An increase in the cutoff correlates with a gradual augmentation in precision, albeit accompanied by a decrease in the total number of retained CIVs. To optimize the selection of LI score cutoff, we fitted a sigmoid function to the precision variation curve against the cutoff,

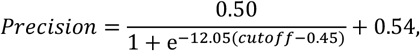

followed by computing the cutoff value that yields a precision of 0.8, which was found to be around 0.6 (0.62). Above this cutoff, there was a strong diminishing return of the increase of precision rate. Therefore, we chose 0.6 as the optimal cutoff and applied it to downstream single-cell genomic data analyses.

### Lineage composition analysis

To better examine the efficacy of CIVs in defining cell lineages in simulation, we developed a method to quantitatively measure the lineage composition of CIV-defined subpopulations. In this method, we first defined the eight cells at the third generation from a common ancestral cell as eight ancestral lineages (termed Lineage 1-8). Then, for each cell within a CIV-defined subpopulation, we traced their ancestor to the third generation (i.e. Lineage 1-8) and calculated the sum of third generation ancestral lineages for each CIV-defined subpopulation. This number serves as an indicator of the performance of CIVs, with values closer to one indicating better performance. For example, if one CIV-defined subpopulation contains ancestors from multiple lineages, the cells within it are not eligible to be considered as one genuine clone, suggesting bad performance of this CIV.

## Acknowledgements

We thank Z. Gu, W. Zhai and Hu laboratory members for constructive discussions. This work was supported by National Natural Sciences Foundation of China (82241236 and 32270693 to Z.H., 32070644 and 32293190 to J.X., 32070870 to Q.M., 32300493 to X.W.), Guangdong Basic and Applied Basic Research Foundation (2021B1515020042 to Z.H.) and China Postdoctoral Science Foundation (2022M723301 to X.W.).

## Author contributions

Z.H., X.W. and K.W. conceived and designed the study. X.W. analyzed the single-cell genomic data and simulated data. K.W. performed simulation studies. W.Z. performed the mtscATAC-seq experiments. Z.T. and D.Z. provided guidance on data analysis and modeling. C.Z. and W.Z.Z. collected samples. Z.H., X.W, J.X. and Q.M. interpreted the results. X.W., Z.H. and K.W. wrote the manuscript with contributions from all co-authors. Z.H, J.X. and Q.M. supervised the project.

## Data availability

The mtscATAC-seq data of two PBMC samples have been deposited in National Genomics Data Center (Nucleic Acids Res 2022), China National Center for Bioinformation (GSA-Human: HRA007291) that are publicly accessible at https://ngdc.cncb.ac.cn/gsa-human. Smart-seq2 dataset of the early human embryo is publicly available at ArrayExpress under accession E-MTAB-9388. Smart-seq2 dataset of T cells is publicly available at GEO under accession GSE193439.

## Code availability

All custom code used to reproduce the analysis is available at GitHub: https://github.com/BoxWong/MT_lineage_tracing.

## Supplementary Information

Supplementary Figs. 1-8

**Supplementary Fig. 1.**
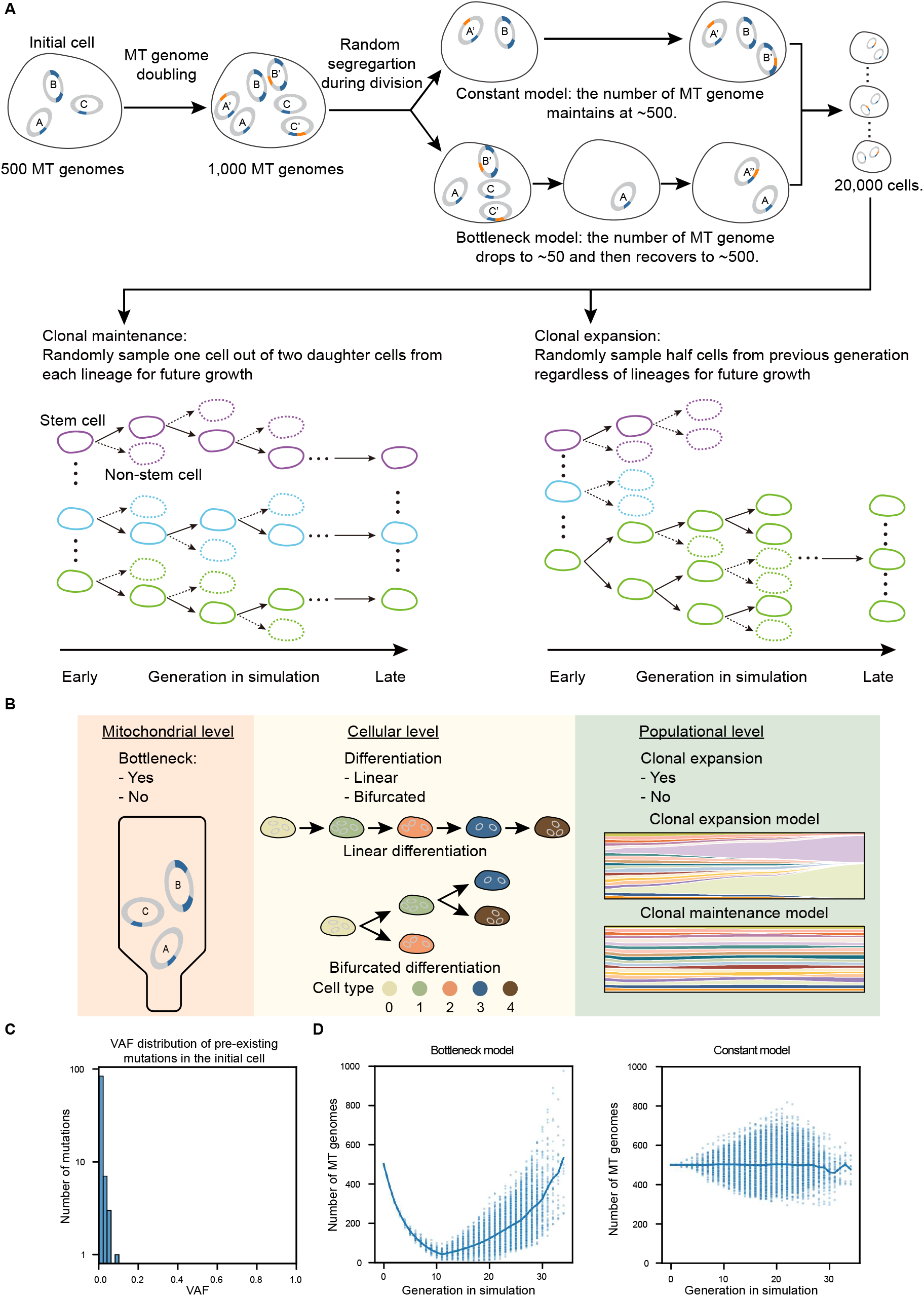
Computational framework of the simulation. **A**, Simulation of mtDNA mutations and transcriptomes for single cells in a dividing population. The first cell was initialized with 500 mitochondrial genomes carrying around 100 pre-existing mutations (blue) whose initial VAFs follow a power-law distribution. During the mitochondrial DNA replication (e.g. A’ is a replicate of A), *de novo* mutations (orange) arose and were accumulated in the cell. After replication of MT genomes, all MT genomes were randomly segregated into two daughter cells. This replication-segregation process was repeated along with the cell divisions. In constant model, MT genome copies remained stable (∼500) along the cell divisions. However, in the bottleneck model, the number of MT genomes first decreased to ∼50 and then recovered back to ∼500. After the recovery of MT genome copies, both constant model and bottleneck model experienced the same replication segregation of MT genomes until the population size reached 20,000. Next, 5,000 randomly sampled cells were subjected to clonal maintenance model or clonal expansion model. For clonal maintenance model, each cell was divided into two and only one cell was kept for future divisions (stem cell asymmetric division), whereas in clonal expansion model, half cells from each generation, regardless of their lineages, were kept for future divisions, resulting strong lineage turnover. During the cell division, transcriptomic profiles (n=4,000 genes) were simulated to follow the linear differentiation or bifurcated differentiation. **B**, Overview of the models in this study. At the mitochondrial level, scenarios with or without mitochondrial bottleneck were simulated. At the cellular level, two differentiation scenarios (linear and bifurcated) were simulated. At the populational level, clonal expansion and clonal maintenance were simulated. In total, eight scenarios were simulated and studied. **C**, The VAF distribution of pre-existing mtDNA mutations in the initial cell. **D**, The number of MT genomes over the generation in bottleneck model (left) or constant model (right).

**Supplementary Fig. 2.**
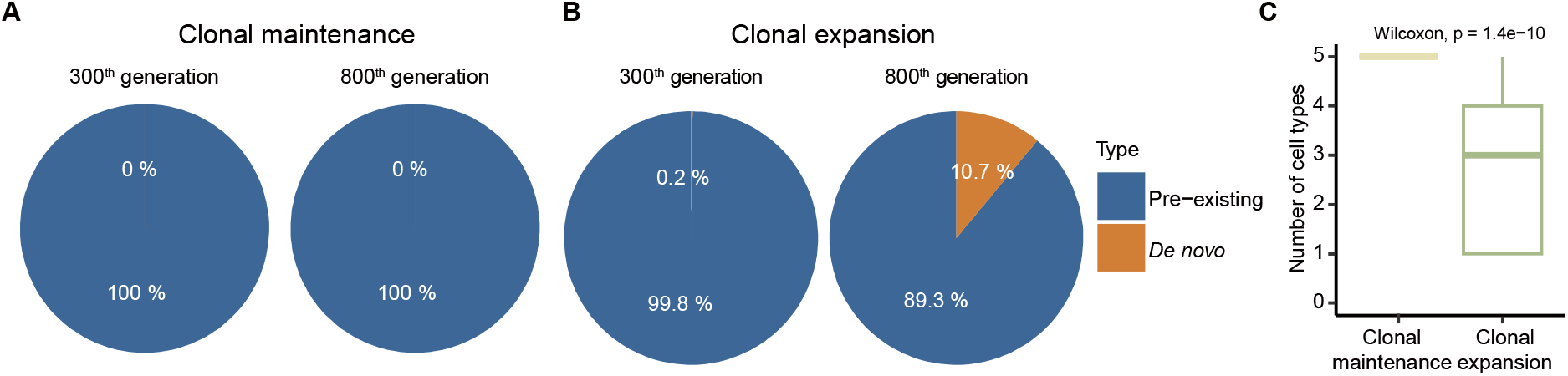
Evaluation of the performance of CIVs in linear differentiation model with mitochondrial bottleneck. **A**, Pie charts showing the proportion of pre-existing and *de novo* mutations of CIVs identified from 300^th^ generation (left) and 800^th^ generation (right) in clonal maintenance model. **B**, Pie charts showing the proportion of pre-existing and *de novo* mutations of CIVs identified from 300^th^ generation (left) and 800^th^ generation (right) in clonal expansion model. **C**, Comparison of cell type composition of each CIV-defined subpopulation between clonal maintenance model and clonal expansion model in linear differentiation with mitochondrial bottleneck.

**Supplementary Fig. 3.**
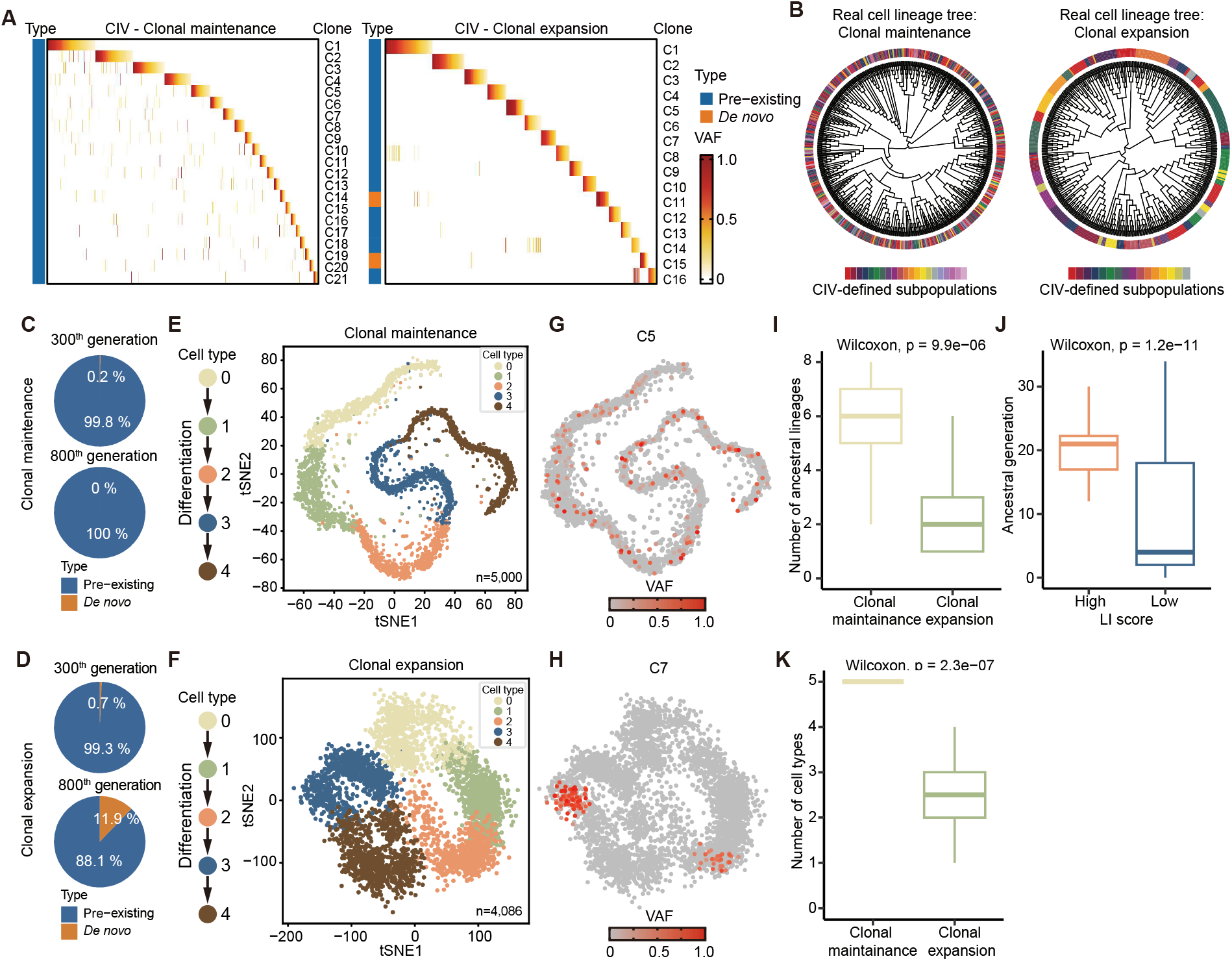
Evaluation of the performance of CIVs in linear differentiation model without mitochondrial bottleneck. **A**, Identification of CIVs in clonal maintenance model (left) or clonal expansion model (right). Each row represents a CIV and each column represents a cell. **B**, Ground-truth cell lineage trees with CIV-defined subpopulations annotated in clonal maintenance (left) or clonal expansion (right) model. **C**, Pie charts showing the proportion of pre-existing and *de novo* mutations of CIVs identified from 300^th^ generation (top) or 800^th^ generation (bottom) in clonal maintenance model. **D**, Pie charts showing the proportion of pre-existing and *de novo* mutations of CIVs identified from 300^th^ generation (top) or 800^th^ generation (bottom) in clonal expansion model. **E**, Simulated transcriptomic profiles of single cells, following linear differentiation (left), in clonal maintenance model. **F**, Simulated transcriptomic profiles of single cells, following linear differentiation (left), in clonal expansion model. **G**, An exemplar CIV (C5) where the defined subpopulations are highlighted in clonal maintenance model. **H**, An exemplar CIV (C7) where the defined subpopulations are highlighted in clonal expansion model. **I**, Comparison of ancestral lineage composition of each CIV-defined subpopulation between clonal maintenance model and clonal expansion model. **J**, Comparison of ancestral generations of CIV-defined subpopulations between high LI score (≥0.6) groups and low LI score (<0.6) groups. **K**, Comparison of cell type composition of each CIV-defined subpopulation between clonal maintenance model and clonal expansion model.

**Supplementary Fig. 4.**
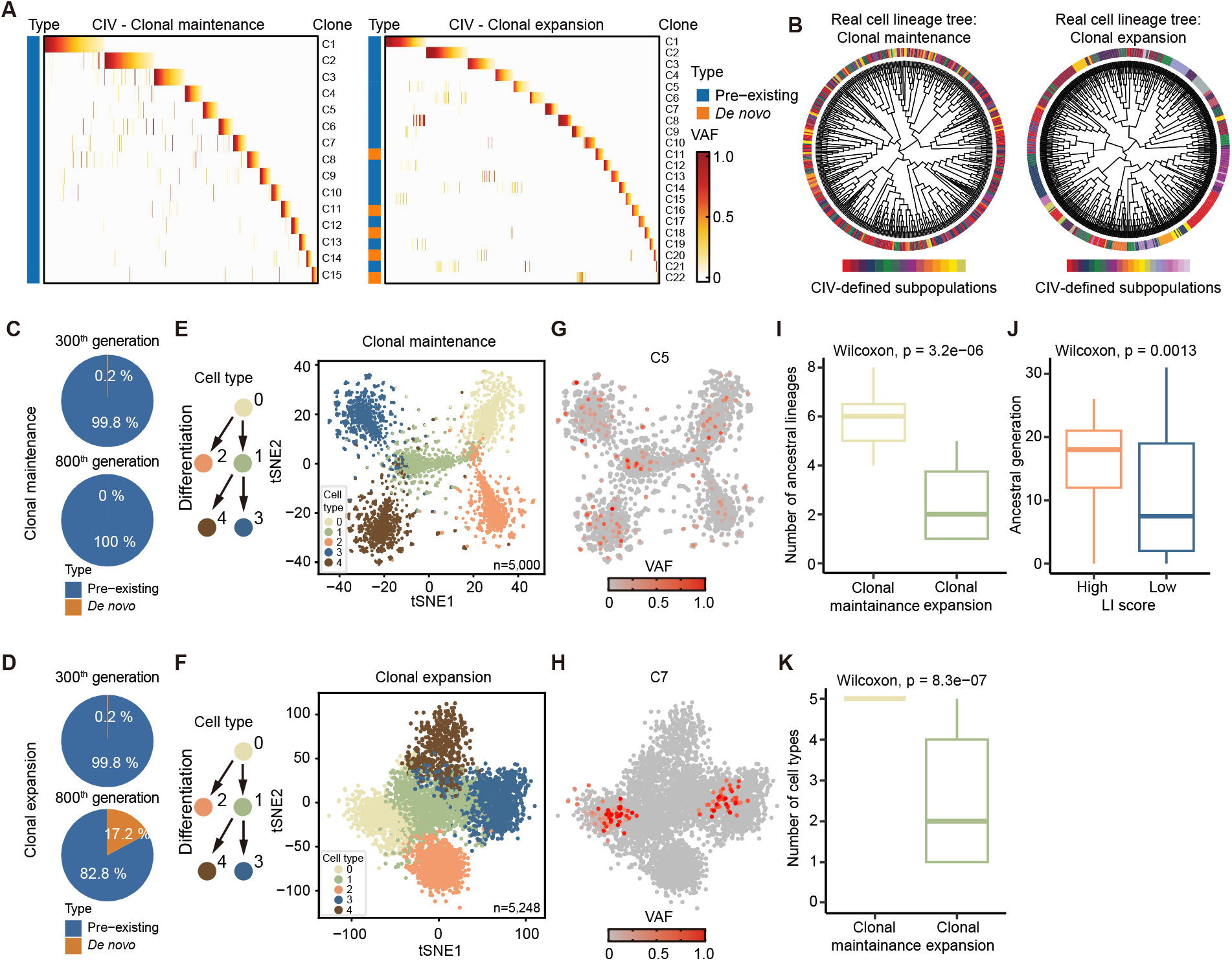
Evaluation of the performance of CIVs in bifurcated differentiation model with mitochondrial bottleneck. **A**, Identification of CIVs in clonal maintenance model (left) and clonal expansion model (right). Each row represents a CIV and each column represents a cell. **B**, Ground-truth cell lineage trees showing the phylogeny of CIV-defined subpopulations in clonal maintenance (left) and clonal expansion (right) model. **C**, Pie charts showing the proportion of pre-existing and *de novo* mutations of CIVs identified from 300^th^ generation (top) and 800^th^ generation (bottom) in clonal maintenance model. **D**, Pie charts showing the proportion of pre existing and *de novo* mutations of CIVs identified from 300^th^ generation (top) and 800^th^ generation (bottom) in clonal expansion model. **E**, Simulated transcriptomic profiles of single cells, following bifurcated differentiation (left), in clonal maintenance model. **F**, Simulated transcriptomic profiles of single cells, following bifurcated differentiation (left), in clonal expansion model. **G**, An exemplar CIV (C5) where the defined subpopulations are highlighted in clonal maintenance model. **H**, An exemplar CIV (C7) where the defined subpopulations are highlighted in clonal expansion model. **I**, Comparison of ancestral lineage composition of each CIV-defined subpopulation between clonal maintenance model and clonal expansion model. **J**, Comparison of ancestral generations of CIV-defined subpopulations between high LI score (≥0.6) groups and low LI score (<0.6) groups. **K**, Comparison of cell type composition of each CIV-defined subpopulation between clonal maintenance model and clonal expansion model.

**Supplementary Fig. 5.**
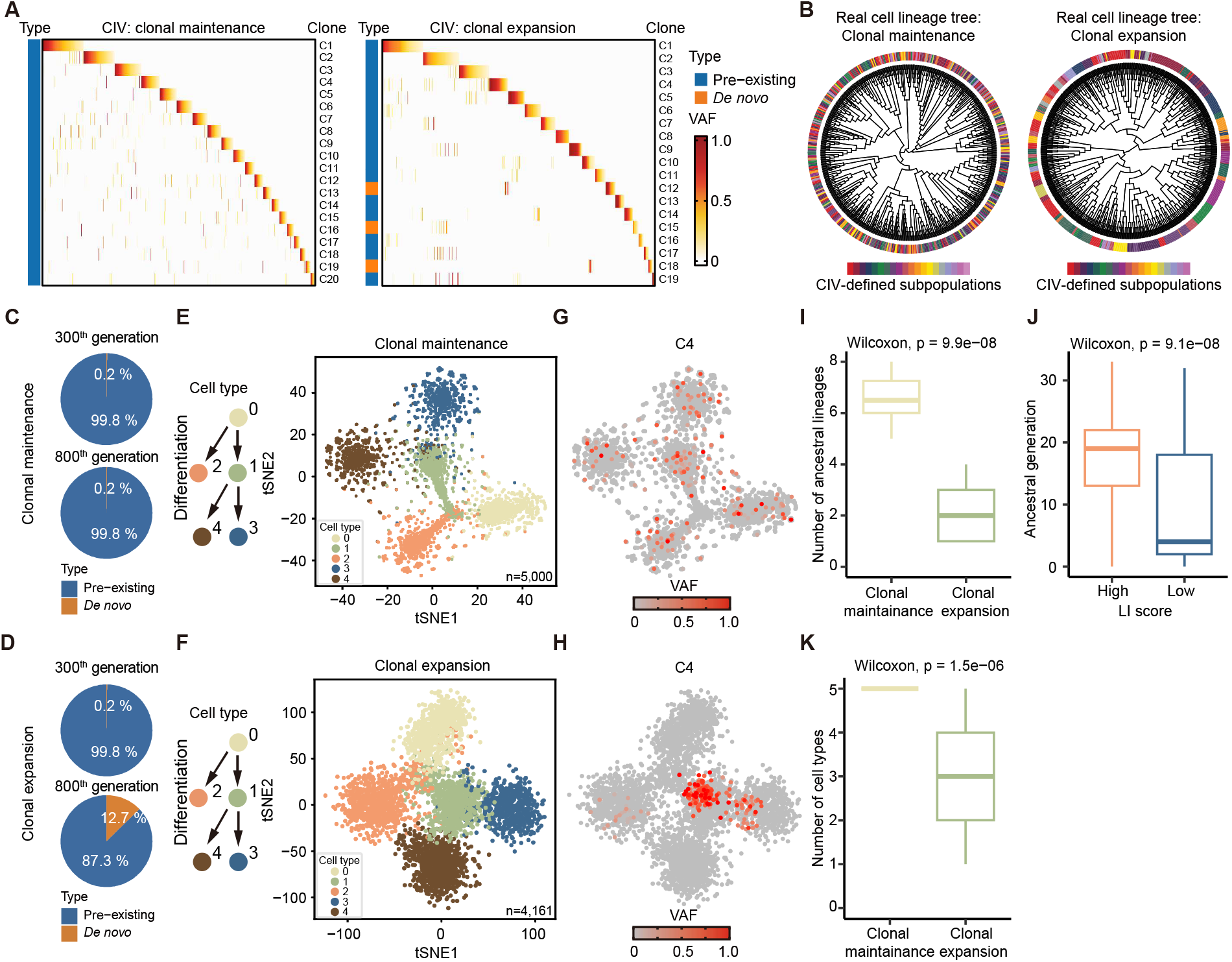
Evaluation of the performance of CIVs in bifurcated differentiation model without mitochondrial bottleneck. **A**, Identification of CIVs in clonal maintenance model (left) and clonal expansion model (right). Each row represents a CIV and each column represents a cell. **B**, Ground-truth cell lineage trees showing the phylogeny of CIV-defined subpopulations in clonal maintenance (left) and clonal expansion (right) model. **C**, Pie charts showing the proportion of pre-existing and *de novo* mutations of CIVs identified from 300^th^ generation (top) and 800^th^ generation (bottom) in clonal maintenance model. **D**, Pie charts showing the proportion of pre-existing and *de novo* mutations of CIVs identified from 300^th^ generation (top) and 800^th^ generation (bottom) in clonal expansion model. **E**, Simulated transcriptomic profiles of single cells, following bifurcated differentiation (left), in clonal maintenance model. **F**, Simulated transcriptomic profiles of single cells, following bifurcated differentiation (left), in clonal expansion model. **G**, An exemplar CIV (C4) where the defined subpopulations are highlighted in clonal maintenance model. **H**, An exemplar CIV (C4) where the defined subpopulations are highlighted in clonal expansion model. I, Comparison of lineage composition of each CIV-defined subpopulation between clonal maintenance model and clonal expansion model. **J**, Comparison of ancestral generations of >80% of CIV-defined subpopulations between high LI score (≥0.6) groups and low LI score (<0.6) groups. **K**, Comparison of cell type composition of each CIV-defined subpopulation between clonal maintenance model and clonal expansion model.

**Supplementary Fig. 6.**
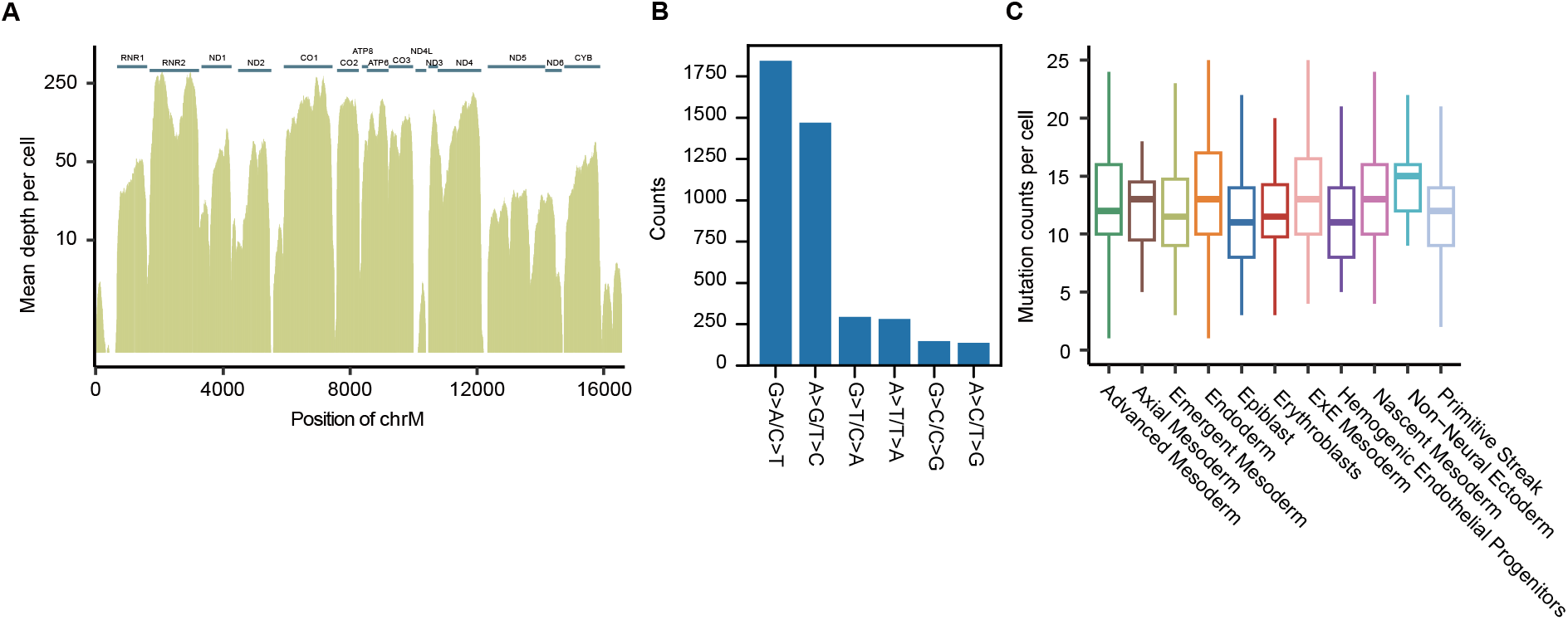
Analysis of Smart-seq2 dataset in an early human embryo. **A**, Sequencing depth of mitochondrial genome per cell in the early human embryo. Lines at the top represent mitochondrial genes. **B**, Mutational pattern of detected mitochondrial DNA mutations in the human embryo. **C**, Box plot showing the mitochondrial DNA mutation counts per cell across cell types.

**Supplementary Fig. 7.**
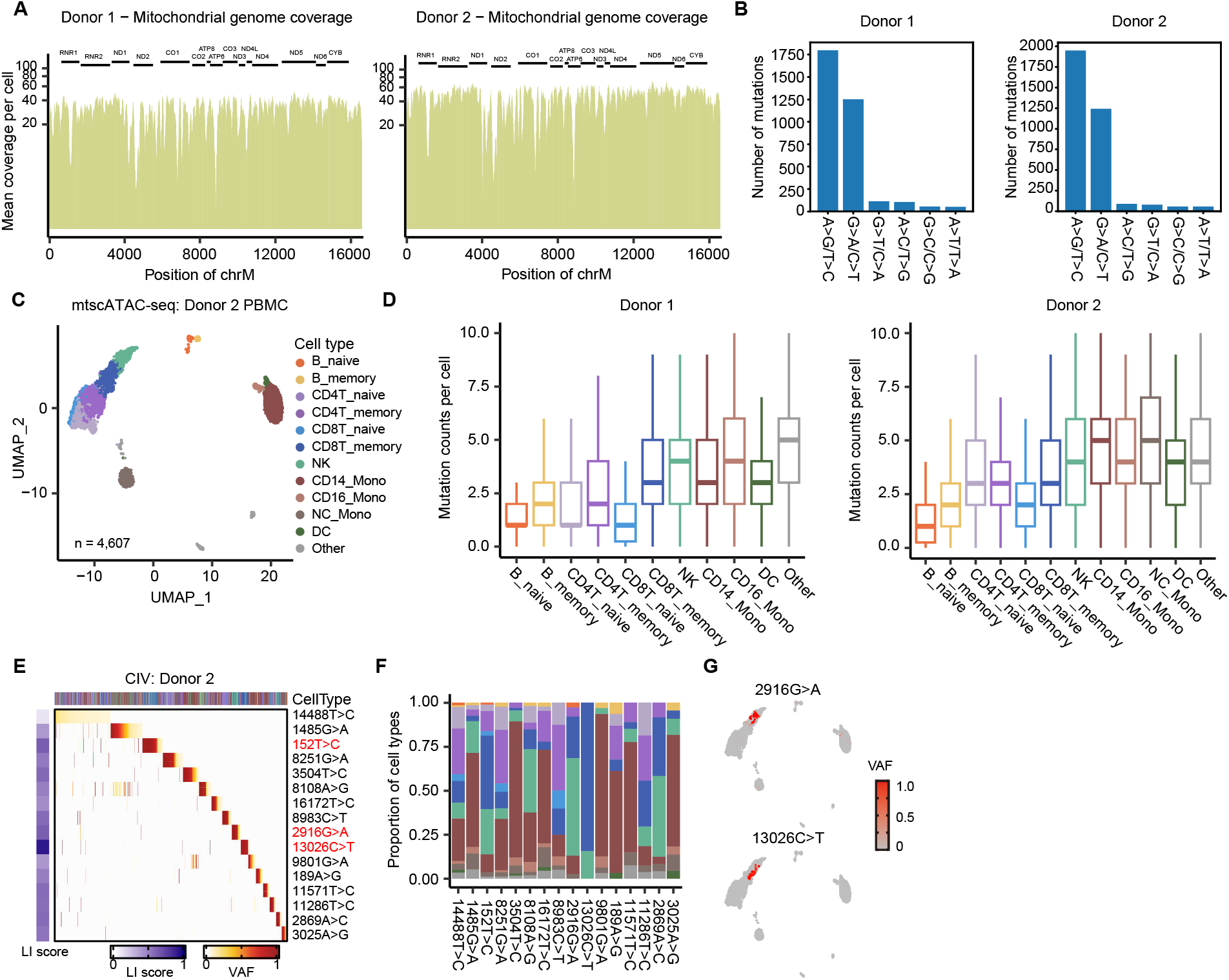
Analysis of mtscATAC-seq dataset of PBMC samples. **A**, Sequencing depth of mitochondrial genome per cell in Donor 1 (left) and Donor 2 (right). Lines at the top represent mitochondrial genes. **B**, Mutational pattern of detected mitochondrial DNA mutations in Donor 1 (left) and Donor 2 (right). **C**, Single-cell chromatin accessibility profiles of PBMC drawn from an old individual, Donor 2 (79 years old). **D**, Box plot showing the mitochondrial DNA mutation counts per cell across cell types in Donor 1 (left) and Donor 2 (right). **E**, Heatmap showing the identification of CIVs in Donor 2. CIVs with LI score greater than 0.6 are highlighted in red. **F**, Barplot showing the proportion of different cell types of CIV-defined subpopulations. **G**, Distribution of 2916G>A- (top) and 13026C>T-defined subpopulations (bottom) on the single-cell chromatin accessibility profile.

**Supplementary Fig. 8.**
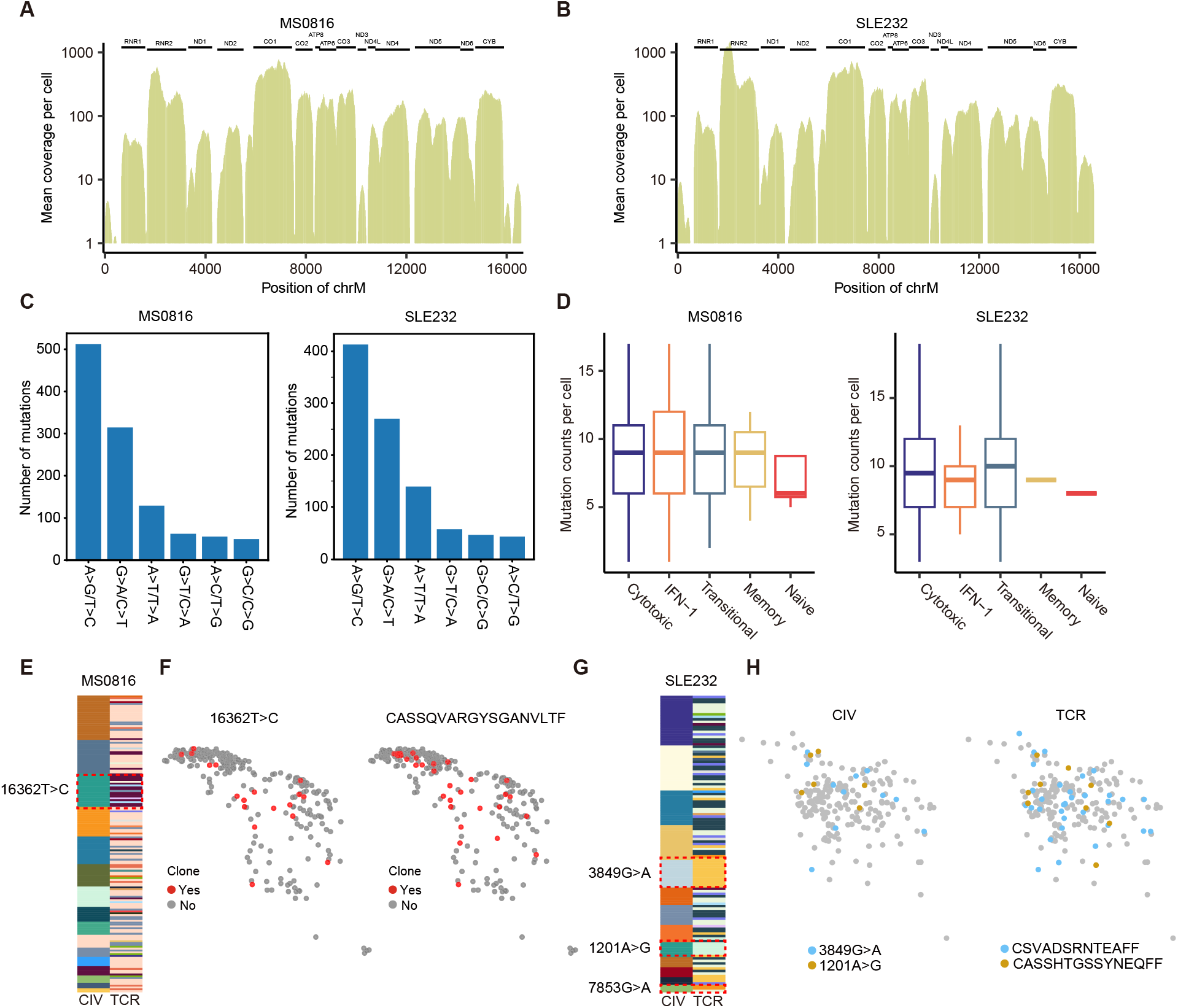
Analysis of smart-seq2 dataset of T cells. **A**, Sequencing depth of mitochondrial genome in a multiple sclerosis (MS) sample - MS0816. Lines at the top represent mitochondrial genes. **B**, Sequencing depth of mitochondrial genome in a systemic lupus erythematosus (SLE) sample - SLE232. Lines at the top represent mitochondrial genes. **C**, Mutational pattern of detected mitochondrial DNA mutations in MS0816 (left) and SLE232 (right). **D**, Box plot showing the mitochondrial DNA mutation counts per cell across T cell subgroups in MS0816 (left) and SLE232 (right). **E**, Correlation of CIV-defined subpopulations and TCR-defined subpopulations in MS0816. Each color represents one subpopulation defined by CIV or TCR. **F**, UMAP diagram showing the distribution of cells within the CIV-defined subpopulation (16362T>C) and the TCR-defined subpopulation (CASSQVARGYSGANVLTF). **G**, Correlation of CIV-defined subpopulations and TCR-defined subpopulations in SLE232. Each color represents one subpopulation defined by CIV or TCR. **H**, UMAP plots showing the distribution of cells within the CIV-defined subpopulations (3849G>A and 1201A>G) and the corresponding TCR-defined subpopulations (CSVADSRNTEAFF and CASSHTGSSYNEQFF).

